# A stress recovery signaling network for enhanced flooding tolerance in *Arabidopsis thaliana*

**DOI:** 10.1101/276519

**Authors:** Elaine Yeung, Hans van Veen, Divya Vashisht, Ana Luiza Sobral Paiva, Maureen Hummel, Bianka Steffens, Anja Steffen-Heins, Margret Sauter, Michel de Vries, Robert Schuurink, Jérémie Bazin, Julia Bailey-Serres, Laurentius A.C.J. Voesenek, Rashmi Sasidharan

## Abstract

Abiotic stresses in plants are often transient and the recovery phase following stress removal is critical. Flooding, a major abiotic stress that negatively impacts plant biodiversity and agriculture, is a sequential stress where tolerance is strongly dependent on viability underwater and during the post-flooding period. Here we show that in *Arabidopsis thaliana* accessions (Bay-0 and Lp2-6), different rates of submergence recovery correlate with submergence tolerance and fecundity. A genome-wide assessment of ribosome-associated transcripts in Bay-0 and Lp2-6 revealed a signaling network regulating recovery processes. Differential recovery between the accessions was related to the activity of three genes: *RESPIRATORY BURST OXIDASE HOMOLOG* (*RBOHD*), *SENESCENCE-ASSOCIATED GENE113* (*SAG113*) and *ORESARA1* (*ORE1/NAC6*) which function in a regulatory network involving a reactive oxygen species (ROS) burst upon de-submergence and the hormones abscisic acid and ethylene. This regulatory module controls ROS homeostasis, stomatal aperture and chlorophyll degradation during submergence recovery. This work uncovers a signaling network that regulates recovery processes following flooding to hasten the return to pre-stress homeostasis.

**Significance statement:** Flooding due to extreme weather events can be highly detrimental to plant development and yield. Speedy recovery following stress removal is an important determinant of tolerance, yet mechanisms regulating this remain largely uncharacterized. We identified a regulatory network in *Arabidopsis thaliana* that controls water loss and senescence to influence recovery from prolonged submergence. Targeted control of the molecular mechanisms facilitating stress recovery identified here can potentially improve performance of crops in flood-prone areas.

## Introduction

Plants continuously adjust their metabolism to modulate growth and development within a highly dynamic and often inhospitable environment. Climate change has exacerbated the severity and unpredictability of environmental conditions that are suboptimal for plant growth and survival, including extremes in the availability of water and temperature. Under these conditions, plant resilience to environmental extremes is determined by acclimation not only to the stress itself, but also to recovery following stress removal. This is especially apparent in plants recovering from flooding. Flooding is an abiotic stress that has seen a recent global surge with dramatic consequences for crop yields and plant biodiversity (1–3). Most terrestrial plants, including nearly all major crops, are sensitive to partial to complete submergence of the above ground organs. Inundations that include aerial organs severely reduces gas diffusion rates, and the ensuing impedance to gas exchange compromises both photosynthesis and respiration. Additionally, muddy floodwaters can almost completely block light access thus further hindering photosynthesis. Ultimately, plants suffer from a carbon and energy crisis and are severely developmentally delayed (4, 5). As floodwaters recede, plant tissues adjusted to the reduced light and oxygen in murky waters are suddenly re-exposed to aerial conditions. The shift to an intensely illuminated and re-oxygenated environment poses additional stresses for the plant, namely oxidative stress and paradoxically, dehydration due to malfunctioning roots, frequently resulting in desiccation of the plant (6). Flooding can thus be viewed as a sequential stress where both the flooding and post-flooding periods pose distinct stressors, and tolerance is determined by the ability to acclimate to both phases.

While plant flooding responses have been extensively studied, less is known about the processes governing the rate of recovery, particularly the stressors, signals, and downstream reactions generated during the post-flood period. When water levels recede, it has been hypothesized that the combination of re-illumination and re-oxygenation triggers a burst of reactive oxygen species (ROS) production. Re-oxygenation has been shown to induce oxidative stress in numerous monocot and dicot species (7–11) and related ROS production dependent on the abundance of ROS scavenging enzymes and antioxidant capacity of tissues (12–16). However, in the link between ROS and survival during recovery, several aspects remain vague, including the source of the ROS and whether it also has a signaling role. Mechanisms regulating shoot dehydration upon recovery also remain to be elucidated. In rice (*Oryza sativa*), the flooding tolerance-associated *SUB1A* gene also confers drought and oxidative stress tolerance during re-oxygenation through increased ROS scavenging and enhanced abscisic acid (ABA) responsiveness (9). In *Arabidopsis*, ABA, ethylene, and jasmonic acid have been implicated in various aspects of post-anoxic recovery (8, 9, 16, 17). While these studies have furthered understanding of flooding recovery, the key recovery signals, the hierarchical relationships between them, and the molecular processes regulating variation and success of recovery remain unclear.

To identify causal mechanisms of the variation in recovery tolerance and unravel the underlying signaling network, we used two *Arabidopsis* accessions Bay-0 and Lp2-6 differing in post-submergence tolerance. The accessions’ sensitivity to complete submergence was primarily due to differences in the shoot tissue during recovery. Through genome-scale sequencing of ribosome-associated transcripts during prolonged submergence and subsequent recovery, we identified three key genes that could explain the superior recovery capacity in Lp2-6: *SENESCENCE-ASSOCIATED GENE113* (*SAG113*), *ORESARA1* (*ORE1/NAC6*), and *RESPIRATORY BURST OXIDASE HOMOLOG* (*RBOHD*). In a network involving a ROS burst, ethylene and ABA, these players regulate ROS homeostasis, stomatal aperture, and senescence to ultimately influence recovery.

## Results

### Submergence Recovery in Two *Arabidopsis* Accessions

*Arabidopsis* accessions Bay-0 and Lp2-6 were previously identified as sensitive and tolerant to complete submergence based on assessment of survival at the end of a recovery period following de-submergence (18). However, further evaluation indicated that this difference in tolerance was mainly due to differences in the recovery phase (Fig. 1*A* and Supplemental Movie). When completely submerged at the 10-leaf stage for 5 days in the dark, plants of both accessions had similar chlorophyll content (Fig. *1B*) and shoot dry weight (Fig. S1*A*). Following return to control growth conditions, however, the tolerant accession Lp2-6 maintained more chlorophyll (Fig. 1*B*) and increased shoot biomass (Fig. S1*A*). Faster development of new leaves in Lp2-6 (Fig. S1*B*) led to higher fitness based on a significantly higher seed yield (Fig. 1*C*). When Bay-0 and Lp2-6 plants were placed in darkness only, rather than submergence together with darkness, both accessions displayed some leaf senescence but no clear phenotypic differences (Fig. S1*C*), indicating that re-aeration determines the distinction in accession survival.

**Fig. 1.**
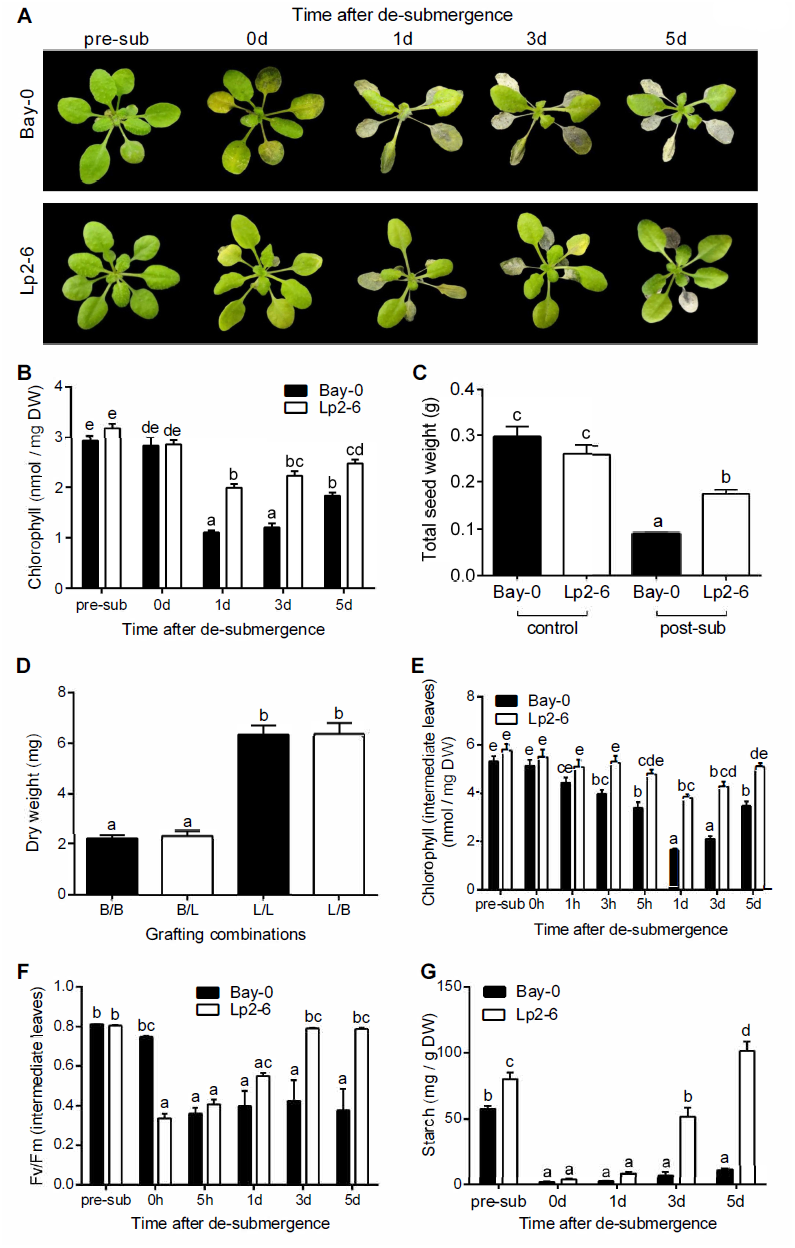
Effects of complete submergence on subsequent recovery in two *Arabidopsis* accessions Bay-0 and Lp2-6. (*A*) Representative shoots of Bay-0 and Lp2-6 before submergence (pre-sub), after 5 d of dark submergence (0 d), and 1, 3, and 5 d of recovery. (*B*) Chlorophyll content of whole rosettes (n=9-10). (*C*) Total seed output of individual control and submergence recovery plants (n=10-15). (*D*) Shoot dry weight of grafted plants submerged for 5 d and recovered for another 5 d under control conditions. Grafting combinations represent the accession of the shoot/root (B=Bay-0; L=Lp2-6) (n=45-60). (*E*) Chlorophyll content in intermediate leaves (n=15). (*F*) Maximum quantum efficiency of photosystem II (Fv/Fm) in intermediate leaves (n=10). (*G*) Starch content in whole rosettes (n=3). Data represent mean ± SEM from independent experiments. Significant difference is denoted by different letters (p<0.05, one-or two-way ANOVA with Tukey’s multiple comparisons test).

The different recovery survival of the accessions was attributed to the shoot since grafting an Lp2-6 shoot to a Bay-0 root or a Lp2-6 root did not affect the high tolerance of Lp2-6 shoots. Similarly, Bay-0 shoots grafted to either Lp2-6 or Bay-0 roots had low tolerance (Fig. 1*D* and Fig. S1*D*). Thus, only shoot traits were further investigated. In both accessions, older leaves showed the most severe submergence damage, with visible dehydration during recovery. Young leaves and the shoot meristem survived in both accessions, but intermediate leaves showed the strongest visible differences between accessions. This correlated with higher chlorophyll content in Lp2-6 intermediate leaves following de-submergence (Fig. 1*E*). Interestingly, photosynthetic capacity after de-submergence, as reflected in Fv/Fm (variable fluorescence/maximal fluorescence), was higher in Bay-0 leaves compared to Lp2-6 leaves (Fig. 1*F*). In subsequent recovery time points, however, Bay-0 intermediate leaves failed to recover towards control Fv/Fm values, whereas Lp2-6 leaves showed full recovery by 3 days following de-submergence. Lower Fv/Fm values in Bay-0 during recovery indicated more photosystem II damage, which may have prevented replenishment of starch reserves (Fig. 1*G*). Based on this characterization of Bay-0 and Lp2-6, further analyses were restricted to the intermediate leaves showing the clearest variable effects of de-submergence stress between both accessions.

### Ribo-seq Reveals Conserved and Accession-Specific Changes in Ribosome-Associated Transcripts During Submergence and Recovery

To identify molecular processes contributing to observed differences in Bay-0 and Lp2-6 during submergence and recovery, the intermediate leaves showing a strong physiological response to de-submergence were subjected to an unbiased ribosome-sequencing (Ribo-seq) approach (18, 19) (Fig. 2 and Fig. S2*A*). Translatome analysis by ribosome footprint sequencing (Ribo-seq) was selected over transcriptome analysis by RNA-seq to increase the likelihood of identifying differentially regulated transcripts that were actively translated, as selective mRNA translation contributes to gene regulation in response to dynamics in oxygen, light, ROS and ethylene (20–24). Intermediary leaves were harvested from plants at the start of the treatment (0h control); submerged in the dark for 5 days (sub) and recovered for 3 hours after de-submergence (rec) (Fig. 2*A*). Each translatome library consisted of at least 38 million reads mapped to the Col-0 genome (Fig. S2*B*). Multidimensional scaling (MDS) showed that biological replicates clustered together (Fig. S2*C*). Furthermore, treatments and accessions clearly clustered separately. Under control conditions the Bay-0 and Lp2-6 translatomes grouped together. As expected, the reads mapped primarily to protein coding regions (Fig. S2*D*).

**Fig. 2.**
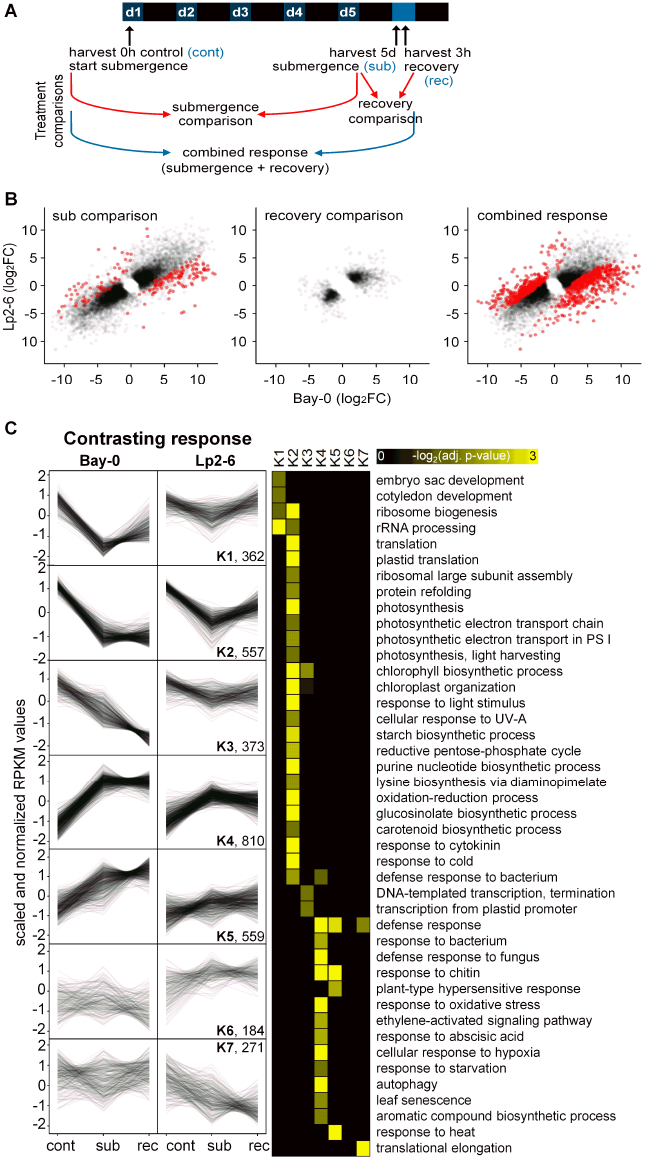
Submergence and recovery induce distinct changes in ribosome-associated transcripts. (*A*) Overview of Ribo-seq experimental design and treatment comparisons. Bay-0 and Lp2-6 intermediate leaves were harvested before treatment (control=cont), 5 d dark submergence (submergence=sub), and 3 h after de-submergence (recovery=rec). The submergence effect was investigated by comparing 5 d submergence-treated samples to the 0 h control (“*submergence comparison*”). Both samples were harvested at the same time during the photoperiod. The recovery effect was a comparison of 5 d submerged samples to those recovered for 3 h in control air and light conditions after de-submergence (“*recovery comparison*”). The combined effect of submergence and recovery was determined by comparing de-submerged 3 h recovery plants with 0 h control plants (“*combined response*”). (*B*) Scatterplots comparing Bay-0 and Lp2-6 log_2_FC under the “*submergence comparison*,” “*recovery comparison*,” and “*combined response*.” Red dots represent accession × treatment DEGs (P_adj_<0.05) and black dots are remaining DEGs. (*C*) Fuzzy K-means clustering of genes showing different behavior in Bay-0 and Lp2-6. Control (0 h, cont), submergence (5 d, sub), and recovery (3 h, rec) conditions were individually plotted as black lines using scaled and normalized RPKM values, and the total number of DEGs in each cluster is noted. GO enrichment for each cluster was visualized as a heatmap.

A large number of genes responded significantly to the treatments and their responses were statistically indistinguishable between the accessions (Fig. 2*B*). These similarly behaving genes were resolved into five clusters using fuzzy K-means clustering (Fig. S3) and enriched gene ontology (GO) categories for these clusters were identified. In both accessions, the common response genes involved in light perception and photosynthesis were downregulated by submergence in darkness but were not re-activated upon recovery (K1). Genes associated with the cytoplasmic translational process were also downregulated (K2), but were upregulated upon recovery. In contrast, responses involved in carbon limitation were strongly induced by submergence and downregulated during recovery (K4). Stress-related GO categories involved in water deprivation and ROS increased upon submergence and rose further during recovery (K5).

To obtain an understanding of processes important for strong performance during recovery, we identified genes at each harvest time-point differing in mRNA abundance between the two accessions (Fig. S2*E*), and genes that responded to the treatments differently (Fig. S2*E* and *F*). Treatment-independent differences increased after submergence and increased even further after the brief recovery period. This was reflected in the number of differentially expressed genes (DEGs) in the accession-specific treatment responses, which was largest when considering the combination of submergence and recovery (Fig. 2*B* and Fig. S2*F*).

Genes with accession-specific regulation were sorted into seven clusters of similarly regulated genes by fuzzy K-means clustering, in which enriched GO categories were identified (Fig. 2*C*). The five largest clusters (K1-K5) of contrasting response genes were characterized by stronger regulation in Bay-0 compared to Lp2-6. During submergence in Bay-0, the GO terms rRNA processing and ribosome biogenesis were strongly downregulated and only marginally recovered upon de-submergence. In Lp2-6, these genes hardly responded to submergence and returned to their original values upon recovery. The same behavior was found in cluster 2 (K2), however, with no recovery in Bay-0 but with a clear recovery response in Lp2-6. GO categories enriched in K2 were related to photosynthesis, light stimuli, and pigment biosynthesis. Cluster 4 (K4), the largest group, was characterized by strong upregulation during submergence and little recovery response in Bay-0. Yet, in Lp2-6, gene induction during submergence was smaller and expression values approached their original control levels during recovery. Corresponding GO categories were related to ethylene and abscisic acid (ABA) signaling, senescence, autophagy, biotic defense, and oxidative stress.

### Inability to Maintain ROS Homeostasis Hinders Recovery

Ribo-seq analyses strongly pointed towards oxidative stress and ROS metabolism as important recovery components. As fuzzy K-means plots revealed, both similarly- and contrastingly-responding genes are overrepresented in GO categories related to oxidative stress (Fig. 2*C* and Fig. S*3*). During submergence, more of these transcripts were associated with ribosomes, with a further increase after 3 hours of recovery. Since this trend was stronger in Bay-0, we investigated the hypothesis that Bay-0 experienced greater oxidative stress thus hindering recovery.

ROS production was measured by assessing levels of the lipid peroxidation product malondialdehyde (MDA). After 5 days of submergence (0 hour after de-submergence), shoot MDA levels were similar to levels in shoots from control non-submerged plants and not different between the accessions (Fig. 3*A*). During subsequent recovery, MDA levels sharply increased in sensitive Bay-0 within 3 hours, and continued to increase over the 3 days of recovery monitored. By contrast, MDA levels in Lp2-6 shoots remained much lower at all recovery time points. ROS production in intermediate leaves was directly quantified using electron paramagnetic resonance (EPR) spectroscopy, which facilitates radical species detection by combination with a spin trapping technique to prolong radical half-life. EPR revealed that ROS content in intermediate leaves under control conditions was close to the detection limit (Fig. 3*B*). Whereas ROS levels were comparable between the accessions at the end of 5 days of submergence, levels began to increase 1 hour after de-submergence in both accessions. This indicated that ROS production is most pronounced following de-submergence. In Bay-0, ROS accumulation peaked at 3 hours of recovery. Afterwards, ROS levels dropped but remained relatively high until the last measurement time point of 24 hours after de-submergence. ROS levels surged in Lp2-6 1 hour after de-submergence, corresponding with concurrent slightly higher MDA production, but subsequently dropped and remained at significantly lower levels than Bay-0 at all subsequent time points. ROS were also measured on intermediate leaves from plants placed in darkness for 5 days followed by recovery in control light conditions (Fig. S4*A*). In both accessions, despite higher ROS levels than control leaves after 5 days of darkness, there was no increase in the recovery period and ROS decreased to the same levels as control plants at 7 and 24 hours of re-illumination. Thus, the ROS burst and ROS content differences during recovery between the two accessions following de-submergence are linked to re-oxygenation rather than reillumination.

**Fig. 3.**
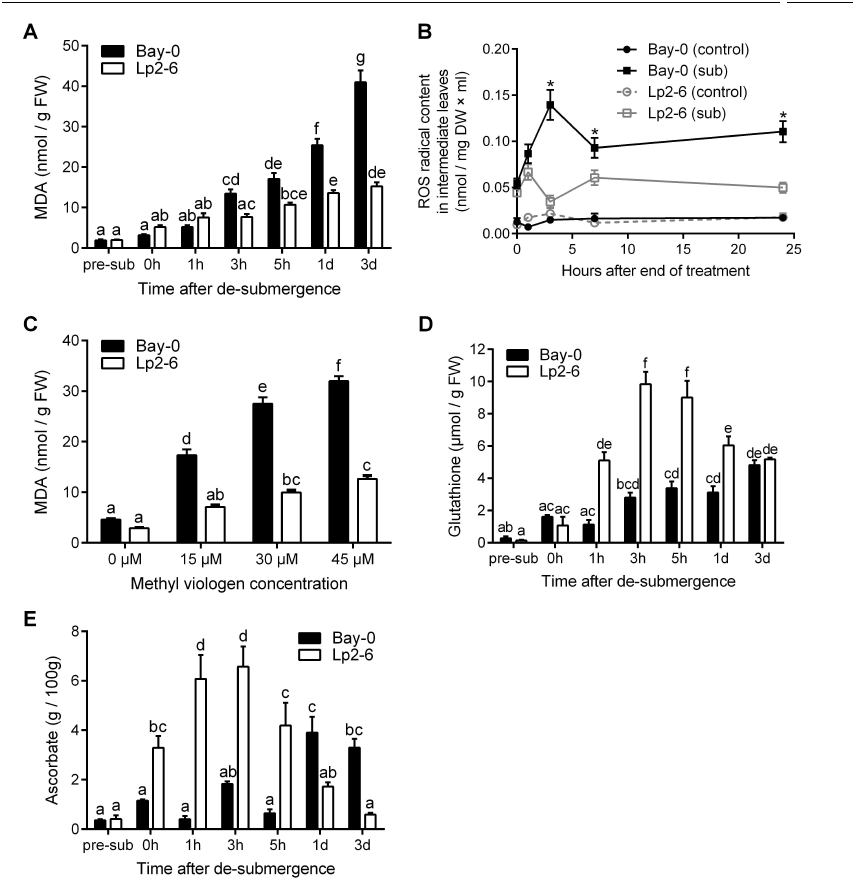
Lp2-6 effectively contains oxidative stress resulting from excessive ROS during recovery. (*A*) Malondialdehyde (MDA) content of Bay-0 and Lp2-6 rosettes before submergence (pre-sub), after 5 d of submergence (0 h), and during subsequent recovery (n=7). (*B*) Electron paramagnetic resonance (EPR) spectroscopy quantified ROS in Bay-0 and Lp2-6 intermediate leaves of control or recovering plants after 5 d of submergence (n=30). Asterisks represent significant difference (p<0.05) between submerged accessions at the specified time point. (*C*) MDA content of rosettes with varying concentrations of exogenously applied methyl viologen (n=7). (*D*) Glutathione and (*E*) ascorbate content in intermediate leaves recovering from 5 d of submergence (n=3). Data represent mean ± SEM. In all panels, except B, significant difference is denoted by different letters (p<0.05, one-or two-way ANOVA with Tukey’s multiple comparisons test).

The direct ROS measurements confirmed that recovery triggered greater ROS accumulation and associated damage in Bay-0. We therefore hypothesized that improved recovery in Lp2-6 is associated with higher oxidative stress tolerance. To assess this, non-submerged plants were sprayed with increasing concentrations of ROS generating methyl viologen (25, 26). For all methyl viologen concentrations tested, Bay-0 had significantly higher MDA levels than Lp2-6, indicating higher ROS-mediated damage and sensitivity to oxidative stress (Fig. 3*C*). To determine whether higher oxidative stress tolerance of Lp2-6 could be a consequence of better ROS amelioration capacity, the antioxidants glutathione and ascorbate were quantified in intermediate leaves. After 5 days of submergence, ascorbate content was significantly higher in Lp2-6, but glutathione levels were similar to that of non-stressed plants in both accessions (Fig. 3*D* and *E*). Starting from 1 hour of recovery, both glutathione and ascorbate increased significantly in Lp2-6, and continued to increase compared to controls (pre-sub) up to 3 to 5 hours after de-submergence. Although ascorbate levels increased in Bay-0, it was delayed compared to Lp2-6 (from 1 day of recovery onwards).

Additionally, we looked for candidate accession-specific genes in the Ribo-seq dataset that could explain higher ROS production in Bay-0. We identified the plasma membrane bound NADPH oxidase *RESPIRATORY BURST OXIDASE HOMOLOGUE* (*RBOHD*; At5g46910) that catalyzes ROS production. Ribosome-associated transcript abundance of *RBOHD* increased during submergence in Bay-0, and recovery conditions further elevated *RBOHD* transcript abundance compared to a moderate induction in Lp2-6 (Fig. S2*D*). This was further confirmed at the level of total transcript abundance by qRT-PCR in an independent experiment (Fig. S4*B*).

To assess the physiological role of *RBOHD* and an associated ROS burst during recovery, the well characterized *rbohD*-*3* loss-of-function mutant (27, 28) was investigated in comparison to its wild-type background Col-0, which is of intermediate submergence tolerance (29, 30). The *rbohD-3* mutant effectively limited ROS production during recovery as discerned by extremely low MDA content in contrast to wild-type Col-0 plants (Fig. *4A* and Fig. S4*A*). However, despite the high MDA content (Fig. 4*A*), wild-type plants recovered from submergence better than *rbohD-3* as reflected in higher chlorophyll content (Fig. 4*B*) and faster new leaf formation (Fig. 4*C;* Fig. S4*C*).

**Fig. 4.**
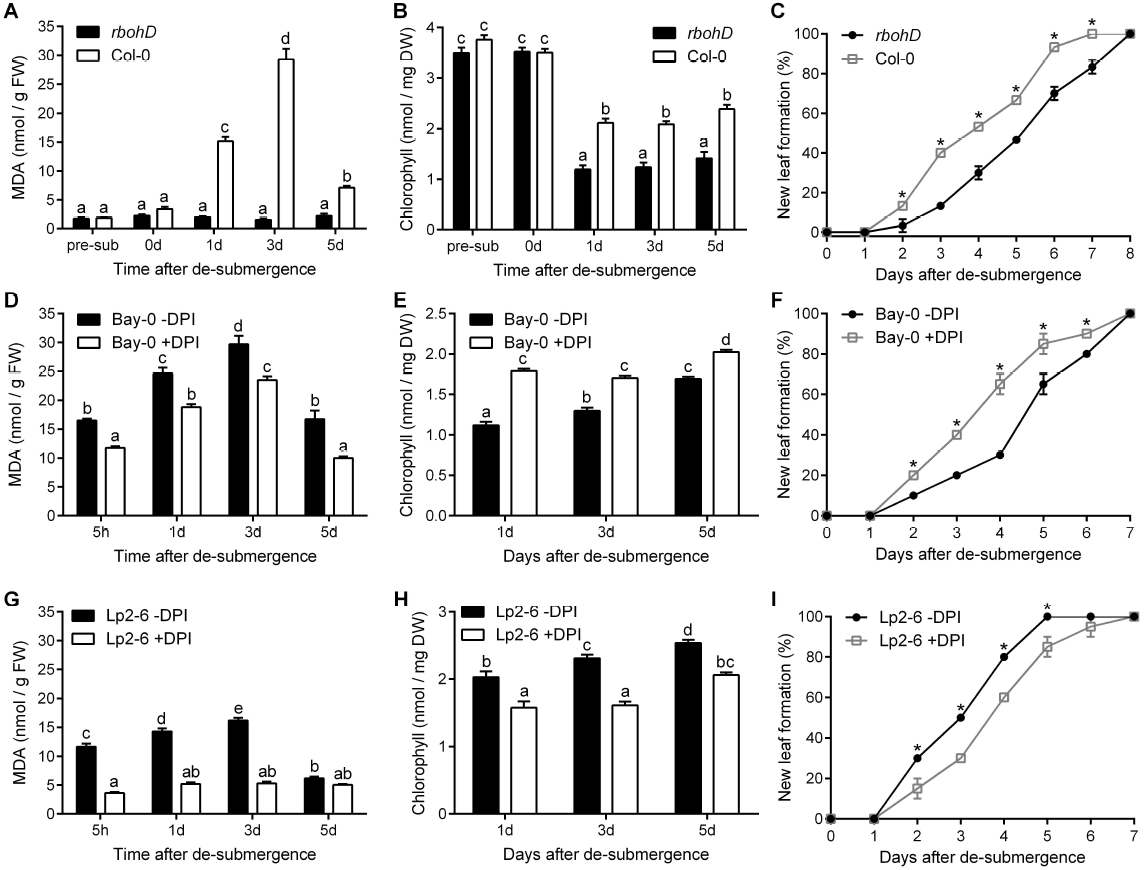
Post-submergence ROS formation mediated through *RBOHD* regulates recovery. (*A*) MDA content (n=12), (*B*) chlorophyll content (n=12), and (C) new leaf formation of *rbohD* and Col-0 (n=30) rosettes during recovery following 5 d of submergence. (*D*) MDA content (n=20), (*E*) chlorophyll content (n=20) and (*F*) new leaf formation (n=20) of Bay-0 plants with or without diphenyleneiodonium (DPI) application upon de-submergence. (*G*) MDA content (n=20), (*H*) chlorophyll content (n=20) and (*I*) new leaf formation during recovery of Lp2-6 plants sprayed with or without DPI upon de-submergence (n=20). Data represent mean ± SEM. Asterisks represent a significant difference between the two accessions at the specified time point (p<0.05, two-way ANOVA with Sidak’s multiple comparisons test). Significant difference is denoted by different letters or (p<0.05, two-way ANOVA with Tukey’s multiple comparisons test).

The necessity of a transient ROS burst involving RBOHD upon de-submergence to initiate signaling might explain slower recovery of *rbohD-3* mutants. However, based on higher *RBOHD* transcript accumulation in Bay-0, we hypothesized that excessive and prolonged ROS production hinders recovery. To test this, the transient ROS burst observed upon de-submergence (3 hours and 1 hour after de-submergence in Bay-0 and Lp2-6 respectively), was manipulated by chemical inhibition of *RBOH* activity. Rosettes were sprayed with the NADPH oxidase inhibitor diphenyleneiodonium (DPI) during the first hour after de-submergence. In Bay-0, DPI application significantly reduced MDA content during recovery (Fig. 4*D*). Furthermore, DPI boosted Bay-0 recovery compared to mock-sprayed plants (Fig. S4*D*), as reflected in significantly higher chlorophyll content within 1 day of recovery (Fig. 4*E*) and faster new leaf development (Fig. 4*F*). For Lp2-6, which accumulated less ROS upon recovery, DPI application further reduced ROS production as indicated by MDA content (Fig. 4*G*). MDA content in DPI-sprayed plants was low at all recovery time points, although slightly higher than levels in *rbohD-3*, whereas mock-sprayed plants had strong MDA accumulation up to 3 days of de-submergence. Even though the dampening of recovery by DPI on Lp2-6 was not as severe as in the *rbohD-3* (Fig. S4*E*), recovery was hindered in DPI-sprayed Lp2-6 plants as indicated by lower chlorophyll content (Fig. 4*H*) and delayed production of new leaves (Fig. 4*I*).

These data demonstrate that excessive ROS accumulation limits recovery, whereas limited and controlled ROS production soon after de-submergence is beneficial for recovery. In Bay-0, DPI application likely dampened the otherwise excessive ROS formed upon de-submergence, thus improving recovery. However, Lp2-6 recovery was hampered when ROS levels were significantly reduced over the recovery time course. We conclude that a fine-tuned balance between production and scavenging of ROS generated by RBOHD and possibly other NADPH oxidases is critical for recovery of leaf formation and ultimately fecundity following de-submergence.

### Dehydration Stress Upon De-Submergence Hampers Recovery

Accession-specific DEGs were also enriched for GO categories associated with dehydration: ABA response and senescence (Fig. 2*C*). Dehydration and senescence were clearly visible during recovery and these symptoms were more severe in Bay-0 (Fig. 1*A*). To assess leaf water management during recovery, relative water content (RWC) was measured in intermediate leaves following de-submergence (Fig. 5*A*). RWC dropped significantly in both accessions 3 hours after de-submergence, although Lp2-6 retained higher water status. RWC values above 70% were maintained at subsequent time points by Lp2-6, while values dropped below 65% by 3 hours and did not recover in Bay-0. A similar trend was observed in water loss assays in detached de-submerged shoots over a 6-hour period. In both accessions, in the first hour after separation from the root, a steep increase in water loss was observed in detached shoots (Fig. 5*B*). Yet, water loss at all subsequent time points was significantly lower in Lp2-6.

**Fig. 5.**
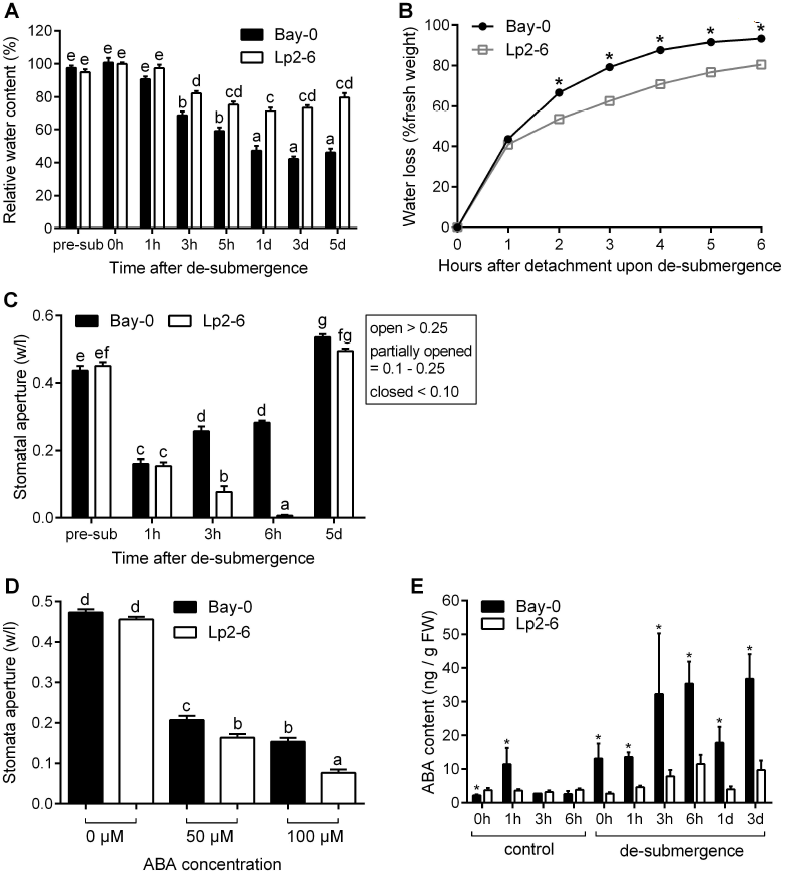
Higher desiccation stress in Bay-0 corresponds with earlier stomatal opening during recovery. (*A*) Relative water content in intermediate leaves before submergence (pre-sub), after 5 d of submergence (0 h), and subsequent recovery time points (n=15). (*B*) Hourly water loss of 10-leaf stage rosettes after detachment from roots immediately upon de-submergence (0 h) compared to the initial fresh weight (n=30). (*C*) Stomatal width aperture (based on width/length ratio) measured using stomatal imprints on the adaxial side of intermediate leaves (n=85-227) of plants before treatment (pre-sub), after 5 d of submergence (0 h), and subsequent recovery time points. (*D*) Stomatal aperture of epidermal peels from intermediate leaves of plants grown under control conditions and incubated in 0, 50, or 100 μM ABA (n=180). (*E*) ABA quantification in intermediate leaves of Bay-0 and Lp2-6 recovering from 5 d of submergence and corresponding controls (n=3). Data represent mean ± SEM. Different letters represent significant difference and asterisks represent significant differences between the accessions at the specified time point (p<0.05, (*B*) two-way ANOVA with Tukey’s multiple comparisons test, (*E*) one-way ANOVA with planned comparisons on log-transformed data).

As rate of water loss is closely linked to stomatal conductance, we investigated whether the differences in dehydration response between the accessions was related to stomatal traits. Stomatal size and density were not significantly different between the two accessions (Fig. S1*E* and Fig. S1*F*). However, stomatal aperture following de-submergence differed between Bay-0 and Lp2-6. While most stomata were partially open in both accessions an hour after de-submergence (Fig. 5*C*), stomatal aperture values further decreased in Lp2-6 and remained low up to 6 hours after de-submergence, indicating stomatal closure. By contrast, Bay-0 stomata reopened by 3 hours and remained open at 6 hours after de-submergence, as indicated by higher stomatal aperture values.

Stomatal aperture regulation in response to drought signals is primarily controlled by ABA, supported by appearance of the “response to ABA” GO category (Fig. 2*C* and Fig. S*3*). To examine stomatal responsiveness to exogenous ABA in the two accessions, abaxial epidermal peels from non-stressed plants were incubated in varying ABA concentrations (Fig. 5*D*). Lp2-6 was more sensitive to ABA, with significantly smaller stomatal apertures under 50 and 100 μM ABA compared to Bay-0. To determine if differences in ABA content contributed to the contrasting stomatal aperture response in Bay-0 and Lp2-6, ABA levels were measured in intermediate leaves after de-submergence and during the corresponding circadian light time points (Fig. 5*E*). Average ABA content in Bay-0 was higher after 5 days of submergence (0 hour of de-submergence) and at all subsequent recovery time points up to 3 days of recovery. Since the ABA measurements did not reconcile with the role of ABA as a positive regulator of stomatal closure, we explored the data for de-submergence-associated signals that might antagonize ABA action.

### Ethylene Accelerates Dehydration and Senescence During Recovery in Bay-0 Mediated by *SAG113* and *ORE1*

The Ribo-seq data revealed accession-specific genes in the “ethylene-activated signaling pathway” (Fig. 2*C*). To further investigate the role of ethylene in the differential submergence recovery responses of the two accessions, whole plant ethylene emission was measured. Ethylene production was significantly higher in Bay-0 than in Lp2-6 after 5 days of submergence (0 hours of de-submergence), and this trend persisted 1 hour and 1 day after de-submergence (Fig. 6*A*). Ethylene production in Lp2-6 was almost half that in Bay-0. To investigate whether this ethylene was causal to the stomatal response and plant performance, as reflected in higher chlorophyll loss in Bay-0 during recovery, ethylene action was blocked using 1-methylcyclopropene (1-MCP). Treatment of Bay-0 plants with 1-MCP following de-submergence strongly reduced the number of open stomata (Fig. 6*B*) and decline in chlorophyll content (Fig. 6*C*). We next explored the Ribo-seq dataset for genes that might mediate the ethylene effect on stomatal behavior and chlorophyll loss during recovery. Amongst the accession-specific genes, we identified two previously confirmed targets of the transcription factor EIN3, a positive regulator of ethylene signaling (31, 32): *SENESCENCE ASSOCIATED GENE 113* (*SAG113*; At5g59220) and the transcription factor *NAC DOMAIN CONTAINING PROTEIN 6*/*ORE1*/*ORESARA 1* (*ANAC092*/*NAC2*/*NAC6*; At5g39610).

**Fig. 6.**
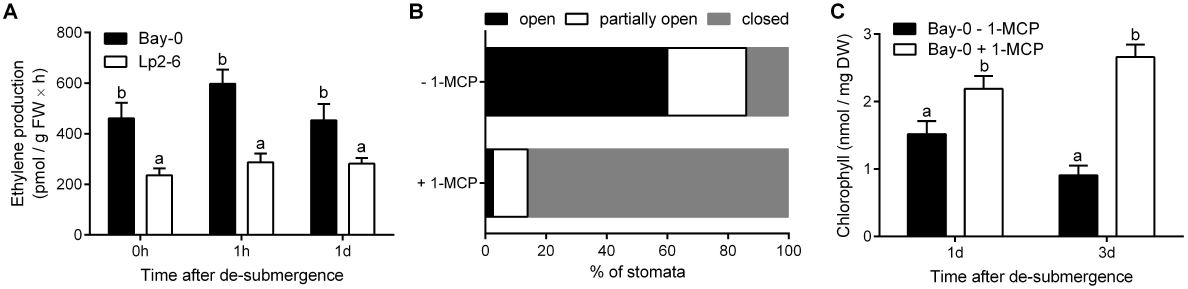
Dehydration and accelerated senescence in Bay-0 upon de-submergence is linked to higher ethylene evolution during recovery. (*A*) Ethylene emissions from Bay-0 and Lp2-6 shoots after de-submergence (n=4-5). (*B*) Stomatal classification at 3 or 6 h after de-submergence of Bay-0 plants treated with or without the ethylene perception inhibitor 1-MCP (n=280-300). (*C*) Chlorophyll content in whole rosettes of Bay-0 treated with or without 1-MCP (n=5-6). 1-MCP treatment was imposed immediately upon de-submergence. Data represent mean ± SEM. Different letters represent significant difference (p<0.05, two-way ANOVA with Tukey’s multiple comparisons test).

Both *SAG113* and *ORE1* were identified as accession-specific genes with increased ribosome-associated transcript abundance in Bay-0 during submergence and 3 hours of recovery, whereas Lp2-6 showed low induction of these transcripts. This trend was confirmed using qRT-PCR in an independent experiment assessing total *SAG113* and *ORE1* transcript abundance (Fig. 7*A* and *B*). *SAG113* encodes a protein phosphatase 2C implicated in the inhibition of stomatal closure to accelerate water loss and senescence in *Arabidopsis* leaves (33, 34). *ORE1* has been previously characterized as a positive regulator of leaf senescence (35–37). In accordance with their identity as EIN3 targets, 1-MCP treatment of Bay-0 following de-submergence significantly repressed *ORE1* and *SAG113* transcript abundance increase during recovery (Fig. 7*C* and *D*). Although 1-MCP suppressed the de-submergence promoted transcript accumulation, both *ORE1* and *SAG113* are also reported to be ABA inducible (33, 38). However, application of an ABA antagonist (AA1) (39) significantly suppressed the de-submergence-induced increase in transcript abundance of *SAG113* only (Fig. S5). Accordingly, AA1-treated plants had a higher percentage of closed stomata corresponding with the role of *SAG113* in stomatal closure of senescing leaves (Fig. S5*E*). Effectiveness of the ABA inhibitory action of AA1 was confirmed with rescuing ABA-induced inhibition of seed germination (Fig. S5*F*) and dark-induced senescence as described by (39), and qRT-PCR of the ABA-regulated genes *RD29B* and *RD22* (Fig. S5*C* and *D*).

**Fig. 7.**
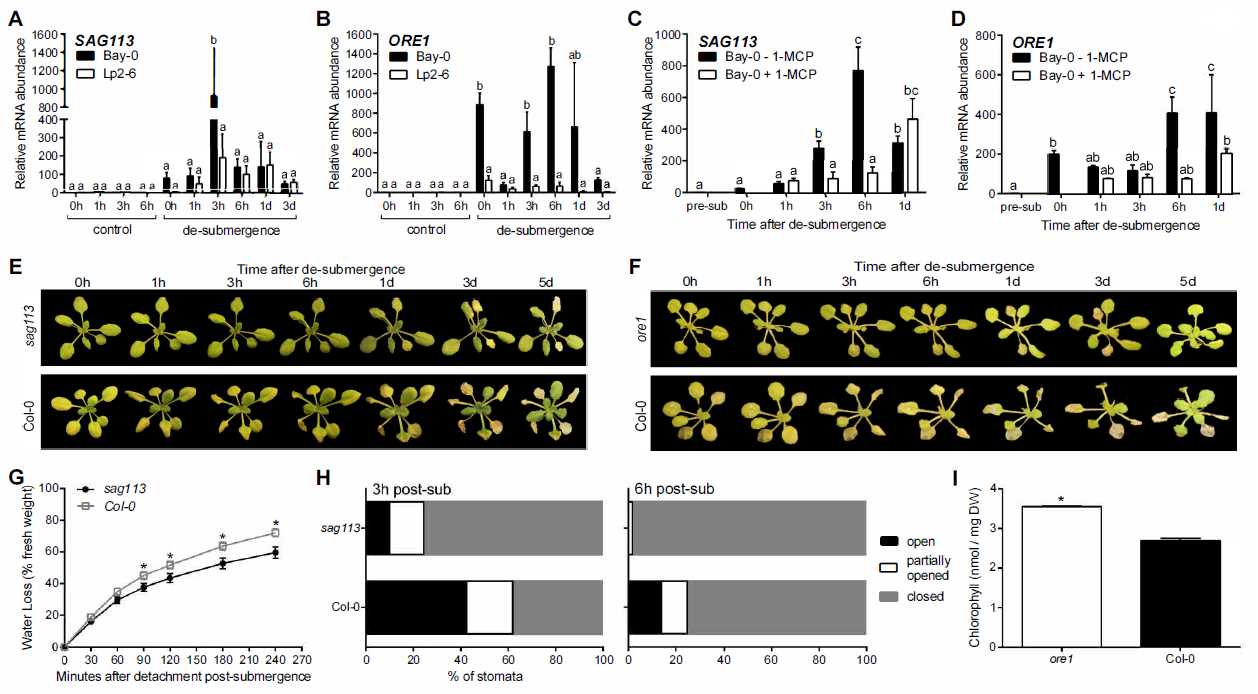
Ethylene-mediated dehydration and senescence in Bay-0 post-submergence links to the induction of *SAG113* inhibiting stomatal closure and *ORE1* promoting chlorophyll breakdown. Relative mRNA abundance of *SAG113* (*A*) and *ORE1* (*B*) measured by qRT-PCR in Bay-0 and Lp2-6 intermediate leaves following de-submergence after 5 d of submergence (n=3 biological replicates). (*C, D*) Relative mRNA abundance of *SAG113* and *ORE1* measured by qRT-PCR in intermediate leaves of Bay-0 plants treated with and without 1-MCP (n=3-4 biological replicates). (*E, F*) Representative images of *sag113* (*E*) and *ore1* (*F*) mutants during recovery after 4 d of submergence compared to wild-type Col-0. (*G*) Stomatal classification at 3 and 6 h after de-submergence for *sag113* and Col-0 submerged for 4 d (n=120-180). (*H*) Water loss in *sag113* and Col-0 after detachment from roots upon de-submergence compared to the initial fresh weight (n=4). (*I*) Chlorophyll content in whole rosettes of *ore1* and Col-0 after 5 d of recovery following 4 d of submergence (n=3). Data represent mean ± SEM. Different letters represent significant difference and asterisks represent significant difference between the genotypes at the specified time point (p<0.05, two-way ANOVA with Tukey’s multiple comparisons test).

Evaluation of a previously characterized knockout mutant for *SAG113* (33, 34), revealed significantly fewer closed stomata at 3 and 6 hours after de-submergence compared to the wild-type Col-0, correlating with significantly reduced water loss (Fig. 7*G* and *H*). Loss-of-function *ore1* mutants (35), had less leaf chlorosis and significantly higher chlorophyll content after 5 days of recovery than in wild-type Col-0 plants (Fig. 7*I*). In conclusion, *SAG113*, induced by the higher ethylene production and ABA levels in Bay-0, contributes to premature stomatal opening and subsequent dehydration. Simultaneously, higher ethylene production in Bay-0 was responsible for *ORE1* induction leading to senescence, as reflected in higher chlorophyll breakdown.

## Discussion

Timely recovery following stress exposure is critical for plant survival. Flooding severely reduces light intensity and gas exchange and subsequent effects on respiration and photosynthesis cause a severe energy and carbon imbalances (2). Floodwater retreat poses new stress conditions as low light- and hypoxia-acclimated plant tissues encounter terrestrial conditions again. Here we exploited two *Arabidopsis* accessions in which differences in submergence tolerance were primarily due to distinctions in submergence recovery. This system revealed that superior recovery after de-submergence is an important aspect of submergence tolerance linked to reproductive output and thus plant fitness (Fig. 1*C*). Using these accessions, we sought to identify molecular and physiological processes and regulatory components influencing recovery.

It is generally accepted that the transition back to re-illuminated and re-oxygenated conditions results in a transient ROS burst in recovering tissues due to reactivation of photosynthetic and mitochondrial electron transport promoting excessive electron and proton leakage (40–42). Reoxygenation led to increased ROS production in both accessions, but sensitive Bay-0 was unable to control prolonged and excessive ROS production during recovery. This could explain the severe photoinhibition (Fig. 1*F*) and hindered starch replenishment in this accession (Fig.1*G)* during submergence recovery. ROS production differences between the two accessions corresponded with higher *RBOHD* transcript abundance during recovery in Bay-0. Counterintuitively, significantly reducing post-submergence ROS generation through genetic (*rbohD-3*) or pharmacological means (DPI application in Lp2-6) worsened recovery. Although excessive ROS are damaging, controlled ROS production via RBOHD might be required for stress signaling during submergence recovery.

ROS production has been previously implicated in hypoxia signaling (43, 44). *RBOHD* is an *Arabidopsis* core hypoxia gene (45, 46) and a transient RBOHD-mediated ROS burst during hypoxia was found to be essential for induction of genes required for hypoxia acclimation (anaerobic metabolism) and seedling survival (44). Pretreatment of *Arabidopsis* seedlings with DPI prior to hypoxia reduced core response gene upregulation and limited survival (43). *RBOHD* is also a candidate gene within a quantitative trait locus conferring submergence tolerance in 10-12 leaf stage *Arabidopsis* (47). Our results demonstrate that *RBOHD* also has an essential role in submergence recovery. In Lp2-6, higher oxidative stress tolerance was linked to restricted ROS accumulation within one hour of de-submergence and a significant increase in antioxidant status (Fig. 3*D* and *E*). Clearly, maintenance of a delicate balance of ROS and antioxidants is critical to cellular homeostasis. While controlled ROS production is essential, it needs to be countered by an effective antioxidant defense system that can manage excessive ROS accumulation and associated damage. The recovery signals regulating *RBOHD* are unclear, but it is likely to be under hormonal control.

Our work also highlighted dehydration stress and accelerated senescence as deterrents to recovery. Plants recovering from flooding often experience physiological drought due to impaired root hydraulics and/or leaf water loss (9, 13, 48, 49). Tolerant Lp2-6 rosettes regulated water loss following de-submergence more effectively than Bay-0. The inferior hydration status of Bay-0 correlated with earlier stomatal re-opening 3 hours following de-submergence. The smaller stomatal apertures of Lp2-6 most probably counteracted dehydration during recovery. The Ribo-seq data and hormone measurements indicated a stronger ABA response in Bay-0, conflicting with the role of ABA in promoting stomatal closure in response to drought signals. However, the Ribo-seq data also revealed a possible role for ethylene signaling in mediating recovery differences between the accessions (Fig. 2*C*). Ethylene is a senescence-promoting hormone that can antagonize ABA action on stomatal closure (50). Elevated ethylene production following de-submergence in Bay-0 corresponded with both an earlier stomatal reopening and greater chlorophyll loss, since chemical inhibition of ethylene signaling during recovery reversed both traits. We suggest that ethylene action is mediated through the EIN3 target genes, *SAG113* and *ORE1*, identified as accession-specific regulated genes with higher transcript abundance in Bay-0 during recovery. Accordingly, knockout mutants in the Col-0 wild-type background, with intermediary submergence tolerance (29), showed improved recovery following de-submergence, associated with improved water loss and reduced senescence. Although previous work on *Arabidopsis* seedlings recovering from anoxic stress (8) revealed that ethylene is beneficial for recovery, our data indicate a negative role for ethylene in submergence recovery. Since ACC conversion to ethylene requires oxygen, ethylene production is limited by anoxic conditions during prolonged submergence (51). Higher ethylene production in Bay-0 upon de-submergence might imply more ACC accumulation during submergence. Upon re-oxygenation, ethylene formation mediated by ACC synthase and oxidase enzymes may accelerate dehydration and senescence by inducing *ORE1* and *SAG113*.

The increase in *SAG113* transcript abundance following de-submergence was reduced upon application of an ABA antagonist indicating ABA regulation of this gene (Fig S5*A*). This implied that high ABA levels in Bay-0 would promote stomatal opening via *SAG113* upregulation, rather than closure, which appears counterintuitive. However, this may reflect interplay between ABA and ethylene signaling pathways. The induction of *SAG113* in Bay-0 could be a means to accelerate senescence of older leaves to remobilize resources to younger leaves, and possibly meristematic regions for new leaf development. How ethylene and ABA interactions influence recovery is an interesting area for future research.

Based on our findings, we propose a signaling network that regulates submergence recovery. Following de-submergence, dehydration caused by reduced root function and re-oxygenation generate the submergence recovery signals ROS, ABA, and ethylene that elicit downstream signaling pathways regulating various aspects of recovery (Fig. 8). Recovery signaling requires *RBOHD*-mediated ROS production, but this must be transient to prevent subsequent oxidative damage and photoinhibition. ABA and ethylene signaling likely interact to control stomatal opening, dehydration and senescence through regulation of genes such as *SAG113* and *ORE1*. This work provides key new insights into the highly regulated processes following de-submergence that limit recovery of Bay-0 and bolster survival of Lp2-6, emphasizing selection on mechanisms enhancing the return to homeostasis.

**Fig. 8.**
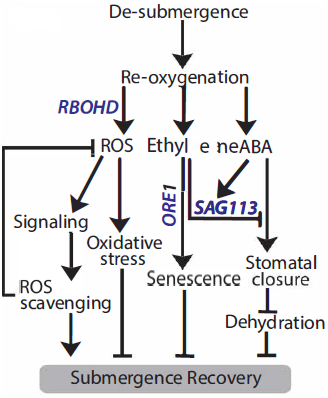
A signaling network mediating post-submergence recovery. Following prolonged submergence, the shift to a normoxic environment generates the post-submergence signals ROS, ethylene, and ABA. A ROS burst upon re-oxygenation occurs due to reduced scavenging and increased production in Bay-0 from several sources, including *RBOHD* activity. While excessive ROS accumulation is detrimental and can cause cellular damage, ROS-mediated signaling is required to trigger downstream processes that benefit recovery, including enhanced antioxidant capacity for ROS homeostasis. Signals triggering *RBOHD* induction following de-submergence are unclear, but hormonal control is most likely involved. Recovering plants experience physiological drought due to reduced root conductance, resulting in increased ABA levels post-submergence which can regulate stomatal movements to offset excessive water loss. High ethylene production in Bay-0 caused by ACC oxidation upon reaeration can counter drought-induced stomatal closure via induction of the protein phosphatase 2C *SAG113*, accelerating water loss and senescence. Higher transcript abundance of *SAG113* in Bay-0 is also positively regulated by ABA and could be a means to speed up water loss and senescence in older leaves. Ethylene also accelerates chlorophyll breakdown via the NAC TF *ORE1*. The timing of stomatal reopening during recovery is critical for balancing water loss with CO_2_ assimilation and is likely regulated by post-submergence ethylene-ABA dynamics and signaling interactions.

## Materials and Methods

### Plant Growth and Submergence Treatment

*Arabidopsis* seeds were obtained from the Nottingham Arabidopsis Stock Centre (NASC, UK) or received from the listed individual: Bay-0 (accession CS22633), Lp2-6 (accession CS22595), Col-0, *rbohD-3* (N9555, containing a single dSpm transposon insertion, received from Ron Mittler, University of North Texas) (27), *sag113* (SALK_142672C, containing a T-DNA insertion) (34), *ore1* (SALK_090154, containing a T-DNA insertion) (35). All mutants were in the Col-0 wild-type background and were genotyped to confirm the presence of the insertion (Table S1). Seeds were sown on a 1:2 part soil:perlite mixture, stratified (4 d in the dark, 4°C), and under short-day light conditions [9 h light, 20°C, 180 μmol m^-2^ s^-1^ photosynthetically active radiation (PAR), 70% relative humidity (RH)]. At 2-leaf stage, seedlings were transplanted into pots with the same soil mixture covered with a mesh. For submergence, disinfected tubs were filled with water for overnight temperature equilibrium to 20°C. Homogeneous 10-leaf stage plants were submerged at 10:00 AM (2 h after the start of the photoperiod) at 20 cm water depth in a dark 20°C temperature-controlled climate room. After 5 d of submergence, de-submerged plants were replaced in normal growth conditions to follow post-submergence recovery.

### Chlorophyll and Dry Weight

Chlorophyll was extracted from whole rosettes or only intermediate leaves with 96% (v/v) DMSO dark-incubated at 65°C and cooled to RT. Absorbance at 664, 647, and 750 nm was measured with a spectrophotometer plate reader (Synergy HT Multi-Detection Microplate Reader; BioTek Instruments Inc., USA). Chlorophyll a and b concentrations were calculated following the equations of (52). Rosettes and leaves were dried in a 70°C oven for 2 d for dry weight measurements.

### Seed Yield

Control and de-submerged plants grown in short-day conditions were watered daily until the terminal bud stopped flowering, and removed from high humidity conditions for drying until all siliques turned brown. Seeds were collected from individual plants and weighed.

### Shoot and Root Grafting

Grafting methods were based on (53). Sterilized seeds sown on ½ MS plates containing 1% (w/v) agar and 0.5% (w/v) sucrose were stratified (3 d in the dark, 4°C) and grown under short-day light conditions for 6 d. Shoots and roots were grafted in a new ½ MS plate and vertically grown for 10 d. Adventitious roots were excised before transplanting seedlings into mesh-covered pots containing 1:2 parts soil:perlite. Plants were grown under short-day conditions until the 10-leaf stage for 5 d dark submergence.

### Chlorophyll Fluorescence (Fv/Fm) Measurements

Fv/Fm was measured in intermediate leaves (leaf 5 of a 10-leaf stage rosette, where leaf 1 is the first true leaf after cotyledon development). Plants were dark-acclimated for 10 min before using a PAM2000 Portable Chlorophyll Fluorometer (Heinz Walz GmbH, Pfullingen, Germany). The sensor was placed at a 5 mm distance from the leaf. Leaves with an Fv/Fm below detection level were marked as dead.

### Starch Quantification

Starch levels were measured in whole rosettes using a commercial starch determination kit (Boehringer, Mannheim, Germany) following the manufacturer’s protocol.

### Ribo-seq Library Construction

Four intermediate leaves of each rosette submerged for 5 d were frozen in liquid nitrogen at 0 h (10:00 AM, immediately upon de-submergence) and 3 h of air-light recovery. Intermediate leaves of 10-leaf stage control plants were harvested at 0 h. 5 mL of packed tissue was used to isolate ribosome-protected fragments. Ribo-seq libraries were prepared following the methods of (54) and (55, 56). Ribo-seq libraries were multiplexed with 2 samples in each lane. Libraries were sequenced with a HiSEQ2500 (Illumina) sequencer with 50 bp single-end reading. Bioinformatic analyses are described in *SI Materials and Methods*.

### Malondialdehyde Measurements

MDA was quantified using a colorimetric method modified from (57). Leaves were pulverized in 80% (v/v) ethanol and the supernatant was mixed with a reactant mixture of 0.65% (w/v) thiobarbituric acid and 20% (w/v) trichloroacetic acid. After 30 min incubation at 95°C, absorbance was measured at 532 and 600 nm with a spectrophotometer plate reader.

### Electron Paramagnetic Resonance (EPR) Spectroscopy

Intermediate leaves were harvested for each treatment (control, dark, and recovery following submergence) and incubated with a TMT-H (1-hydroxy-4-isobutyramido-2,2,6,6-tetramethyl-piperidinium) spin probe. The supernatant was measured on a Bruker Elexsys E500 spectrometer. Further details are listed in *SI Materials and Methods*.

### Methyl Viologen Application

Plants were sprayed methyl viologen (0, 15, 30, 45 μM) containing 0.1% (v/v) Tween-20 1 d before harvesting. Control plants were sprayed with 0.1% (v/v) Tween-20 to account for detergent effects. Plants were sprayed 3 times during the day, each time with 1 mL of solution.

### Antioxidant Measurements

Glutathione was measured with a Promega GSH-Glo Glutathione Assay kit (Madison, USA), following the manufacturer’s procedure using 25-50 mg of fresh tissue. Ascorbate was measured using a kit from Megazyme (K-ASCO 01/14, Wicklow, Ireland), following the microplate assay procedure with 50-75 mg of fresh tissue.

### Scoring New Leaf Development

Leaves were scored as newly formed during recovery from submergence when emergence from the shoot meristem was clearly visible.

### Application of Chemical Inhibitors of *RBOHD*, ABA, and Ethylene

Upon de-submergence, shoots were sprayed with 400 μL of 200 μM DPI (Sigma-Aldrich, St. Louis, USA) containing 0.1% Tween-20 or 100 μM AA1 (C_18_H_23_N_5_OS_2_, product ID: F0544-0152, Life Chemicals Inc., Niagara-on-the-Lake, Canada) containing 0.1% (v/v) DMSO. Control plants were also sprayed with mock solution containing only 0.1% (v/v) Tween-20 or DMSO. Plants were sprayed again with 200 μL of DPI or AA1 30 min and 1 h after the first application. For 1-MCP gassing, plants placed in glass desiccators (22.5 L volume) were gassed with 5 ppm of 1-MCP (Rohm and Haas Company, Philadelphia, USA). Control plants were placed in a separate desiccator to control for humidity effects. After 15 min, plants were replaced in normal growth conditions. 5 ppm of 1-MCP was reapplied to the plants every 4 h during the first day after de-submergence.

### Relative Water Content

Four intermediate leaves per rosette were detached and fresh weight was recorded. Leaves were saturated in water, and saturated weight was measured after 24 h. Leaves were dried in an 80°C oven for 2 d before measuring dry weight. Relative water content was calculated by: [(fresh weight–dry weight)/(saturated weight–dry weight)]×100.

### Rapid Dehydration Assays

Excised rosettes were weighed hourly up to 8 h after cutting and placed in a controlled environment with ambient room temperature (22.3°C, 12 μmol m^-2^ s^-1^ PAR, 63% relative humidity).

### Stomatal Imprints

Adaxial sides of leaves were imprinted using a silicone-based dental impression kit (Coltène/Whaledent PRESIDENT light body ISO 4823; Altstätten, Switzerland). Leaves were gently pressed onto the silicone mixture and removed after solidification. Transparent nail polish was thinly brushed onto the impression and air-dried. Stomata were viewed on the nail polish impression under a Olympus BX50WI microscope (Tokyo, Japan). Stomatal aperture was reported as width (w) divided by length (l) and classified as open (w/l>0.25), partially open (w/l=0.1-0.25), or closed (w/l=0-0.10). Stomatal measurement immediately upon de-submergence after 5 d of submergence was excluded since the mechanical stress of blotting wet leaves forced stomata to open in Lp2-6.

### ABA Treatment in Epidermal Peels

Epidermal peels were obtained from intermediate leaves of 10-leaf stage rosettes 2 h after the light period began. The adaxial side of the leaf was placed on sticky tape, and the petiole was ripped towards the leaf to obtain a transparent film of the abaxial side. Epidermal peels were placed in potassium stomata opening buffer (50 mM KCl + 10 mM MES, pH 6.15) for 3 h under high light (180μmol m^-2^ s^-1^ photosynthetically active radition (PAR)) and incubated for 1 h in stomata closing buffer [2.5 μM CaCl_2_ + 10 mM MES (pH 6.15)] containing 0, 50, or 100 μM ABA. Stomata on the epidermal peels were viewed under a microscope.

### ABA Extraction and Quantification

Intermediate leaves (60-100 mg) were harvested after de-submergence and control samples were harvested at the same time. ABA was extracted as described in (58), quantified by liquid chromatography-mass spectrometry (LC-MS) on a Varian 320 Triple Quad LC-MS/MS. ABA levels were quantified from the peak area of each sample compared with the internal standard, normalized by fresh weight.

### Ethylene Emission Measurements

Ethylene production was measured based on (51). 2 shoots were placed in a 10 mL glass vial and entrapped ethylene was allowed to escape for 2 min before tightly sealing the vials. After 5 h dark incubation, ethylene was collected with a 1 mL injection needle and measured with gas chromatography (Syntech GmbH, Kirchzarten, Germany).

### RNA Extraction and Quantitative Real-Time qPCR

Total RNA was extracted following the Qiagen RNeasy mini kit protocol (Hilden, Germany). For qRT-PCR, single-stranded cDNA was synthesized from 1 μg RNA using random hexamer primers (Invitrogen, Waltham, USA). qRT-PCR was performed on Applied Biosystems ViiA 7 Real-Time PCR System (Thermo Fisher Scientific) with SYBR Green MasterMix (Bio-Rad, Hercules, USA). Primers used are listed in Table S2. Relative transcript abundance was calculated using the comparative 2^-ΔΔCT^ method (59) normalized to *ACTIN2*.

### Data deposition

The data reported in this paper have been deposited in the Sequence Read Archive (SRA) database, https://www.ncbi.nlm.nih.gov/sra (SRA accession: SRP133870, temporary submission ID: SUB3744462).

## Acknowledgments

At Utrecht University, we thank Rob Welschen for managing the growth facilities, Emilie Reinen and Zeguang Liu for genotyping mutant lines, and Yorrit van de Kaa for seed harvesting. We appreciate Timo Staffel at the University of Kiel for assisting with plant growth for EPR measurements. We thank Thomas Girke of University of California, Riverside for guidance on the RiboSeq workflow in R/Bioconductor. This work was supported by grants to R.S. from the Netherlands Organisation for Scientific Research (NWO 016.VIDI.171.006, and NWO-VENI 863.12.013 and grants to J.B.-S. from the US. National Science Foundation (MCB-1021969) and the US Department of Agriculture National Institute of Food and Agriculture Hatch program. E.Y. was supported by a PhD scholarship from Utrecht University.

## Supporting Information

### SI Materials and Methods

#### Dark Treatment

10-leaf stage plants were placed in a dark climate chamber (20°C, 70% relative humidity) with well-maintained watering. After 5 d, plants were replaced in short-day light conditions.

#### Ribo-seq Bioinformatics

Data analysis was performed on a Linux cluster and R using command line tools, Bioconductor R packages, and custom R scripts. Some scripts were adapted from a systemPipeR Bioconductor R package for Ribo-seq experiments (60, 61). Adapters were trimmed from FASTQ files, and reads of 24 to 36 nt were mapped to the *Arabidopsis thaliana* Col-0 genome (TAIR10/Araport11) in combination with the Araport11 annotation (GFF3 file, obtained from araport.org) using the TopHat and Bowtie2 alignment algorithm (version 2.2.5) allowing 2 mismatches. For reads with multiple mapping, reads were first given priority to the transcriptome and also based on alignment quality score. Log_2_ Fold Changes (log_2_FC) and Benjamini-Hochberg-corrected P-values were calculated using Bioconductor R packages “edgeR” and “limma.” Only genes with more than 15 reads in at least one sample were included. First, libraries were normalized for size and compositional bias with TMM normalization (trimmed mean of M-values). A generalized linear model with a full factorial design of treatment (3 levels: control, submergence, and recovery) and accession (2 levels: Bay-0 and Lp2-6) was fitted to the TMM normalized read count data with a negative binomial distribution. Appropriate comparisons of the treatment, accession and interaction coefficients allowed the calculation of log_2_FC and significance for specific treatments, accession-specific treatment responses (accession × treatment interaction) and treatment-independent differences between the accessions. A MDS plot was created with “plotMDS” function within “edgeR” Bioconductor R packages. Samples distance was determined from the top 2000 differing genes in each pairwise comparison. Scatterplots of log_2_FC comparisons were plotted using custom plotting functions on R. Genes behaving similarly and differently in both accessions were separately clustered with fuzzy K-means clustering (R “cluster” library). RPKM values normalized for library composition (TMM, “edgeR”) were scaled so that for each gene, the average RPKM across all samples was zero and standard deviation was one. Scaled RPKM values were used for fuzzy K-means clustering using Euclidean distances metrics and a membership exponent of 1.2. Genes that best represent their cluster over the entire flooding period (Membership Score > 0.5) were used for visual representation of clustering output. These genes were tested for Gene Ontology (GO) enrichment using the “GOseq” Bioconductor package assuming a hypergeometric distribution and Benjamini-Hochberg-corrected P-values.

#### Electron Paramagnetic Resonance (EPR) spectroscopy

3 intermediate leaves were harvested for each treatment (control, dark, and recovery following submergence) and immediately snapped frozen in liquid N_2_ (62). 150 μL of 1 mM TMT-H spin probe dissolved in 1 mM EDTA was added to each sample. Samples were incubated in a 40°C water bath for 90 min. 20 μL of supernatant was drawn up in a capillary tube for measurements on a Bruker Elexsys E500 spectrometer using the “Xepr acquisition and processing suite” software (Bruker Corporation, Billerica, Massachusetts). Measurements were performed at room temperature with the acquisition parameters: modulation frequency 100 kHz, modulation amplitude 1.3 G, receiver gain 60 dB, time constant 81.92 ms, conversion time 40.11 ms, center field 3512.95 G, sweep width 66.8 G, sweep time 41.07 s, and attenuation 30 dB. A calibration curve for the EPR spectrometer measurement was obtained using a nitroxide radical TEMPO (2,2,6,6-tetramethyl-1-piperidinyloxyl). Calculations from double integration of the low field peak yielded the limit of detection as 0.011 mmol/L and the limit of quantification as 0.038 mmol/L. The concentration of TMT radicals was calculated from the area of the double integration of the low-field peak, which was converted into TEMPO radical equivalents using the calibration curve. ROS concentration was calculated based on the TEMPO-equivalents and the Avogadro constant where 1 mol = 6.022 × 10^23^ radicals, normalized by dry weight.

## Supplemental Figure Legends

**Fig. S1.**
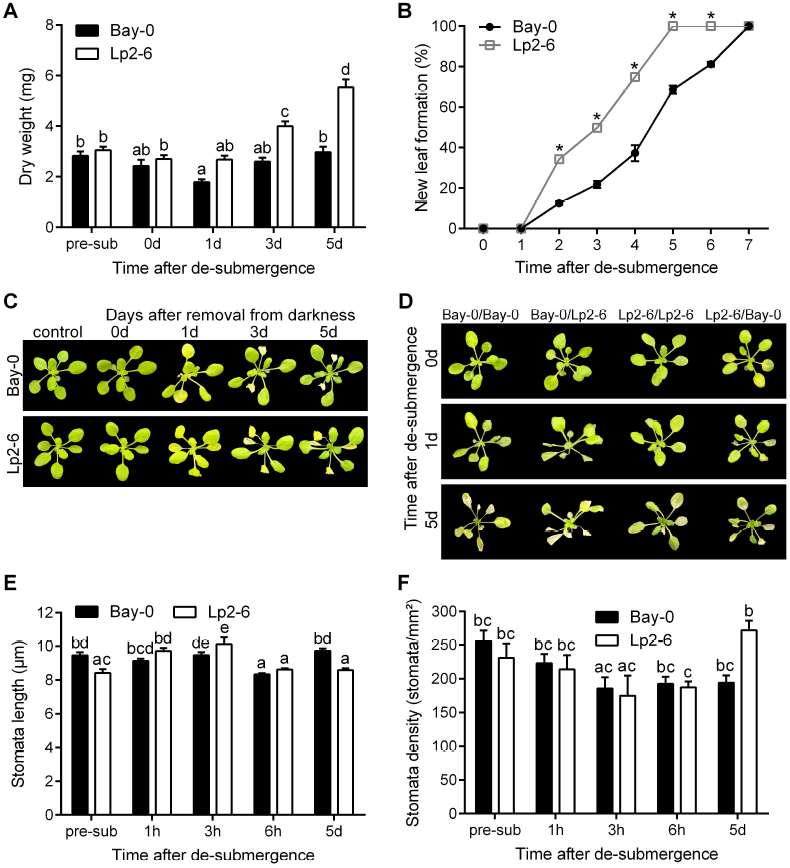
Examining post-submergence recovery using a comparative *Arabidopsis* system. (*A*) Dry weight of whole rosettes during recovery after 5 d of dark submergence (n=9-10). (*B*) Percentage of plants forming new leaves during each day of recovery (n=32). Asterisks represent significant difference between the two accessions at the specified time points (p<0.05, two-way ANOVA). (*C*) Recovery of 10-leaf stage plants after 5 d of darkness as a control for dark submergence. (*D*) Representative images of grafted shoots after 5 d of submergence followed by recovery for 5 d. Images are shown for 0, 1, and 5 d of recovery. Sample groups represent the accession of the shoot/root. (*E*) Stomatal length measured on de-submerged intermediate leaves (n=83-227). (*F*) Stomatal density obtained from abaxial imprints of de-submerged intermediate leaves (n=12-29). Data represent mean ± SEM. Different letters represent significant difference (p<0.05, two-way ANOVA with Tukey’s multiple comparisons test).

**Fig. S2.**
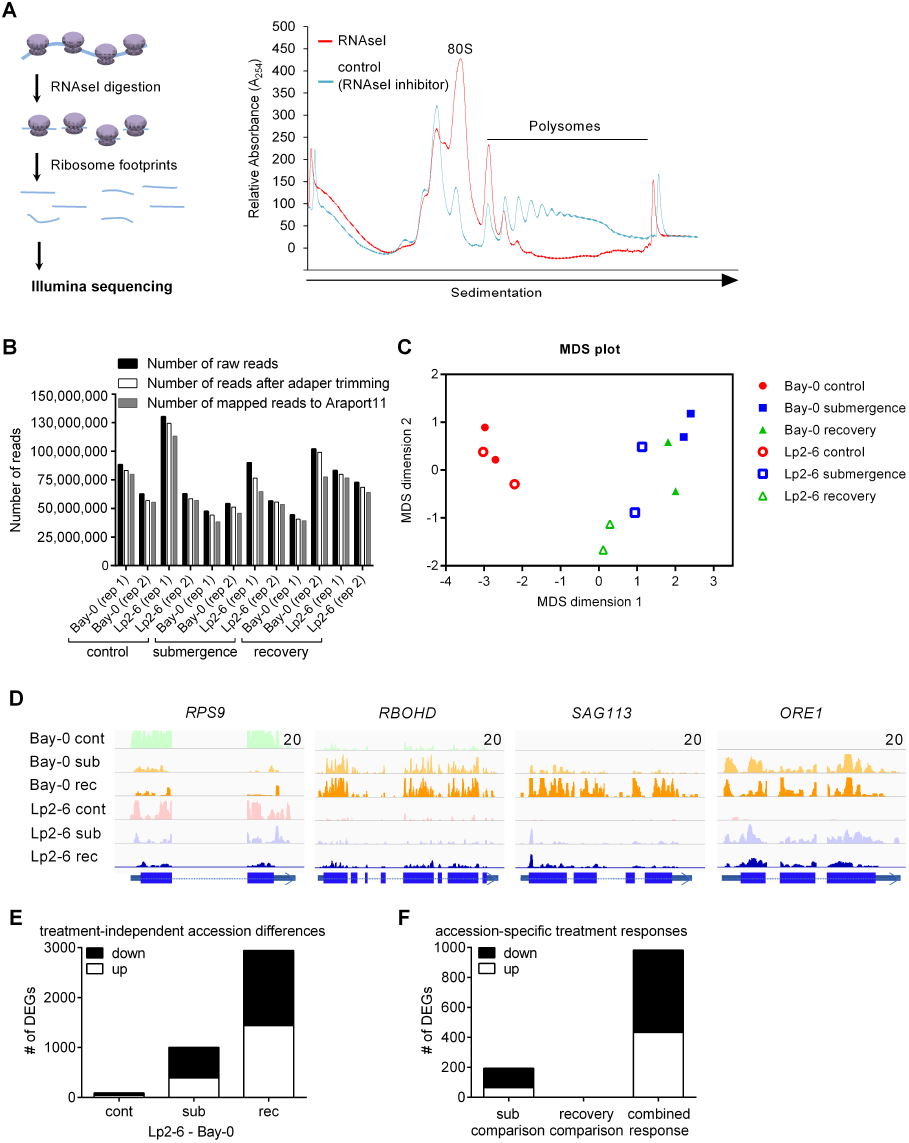
Ribo-seq pipeline for identifying post-submergence molecular mechanisms. (*A*) Representative 254 nm absorbance spectra of sucrose density gradient fractionated control (undigested) and RNAse I digested polysomes. The *x*-axis corresponds to the gradient, with the orientation of sedimentation shown. The first two peaks at the left represent the 40S and 60S monosomes, followed by the 80S peak and the denser polysome peaks. mRNA regions protected by ribosomes from digestion (ribosome footprints) were isolated and constructed into cDNA libraries. (*B*) Illumina sequencing yielded high numbers of reads. Raw reads were unprocessed read output, trimmed reads were those with adapter sequences removed, and mapped reads were those aligning to the Araport11 Col-0 annotated genome. (*C*) Multidimensional scaling (MDS) plot shows distribution of the 2 biological replicates of air control, submergence, and recovery samples. Sample distances were calculated based on the top 2000 pairwise contrasting genes. (*D*) Gene view of coverage of ribosome footprints on 4 genes: nuclear-encoded plastid *RIBOSOMAL PROTEIN S9* (*RPS9*; At1g74970), *RESPIRATORY BURST OXIDASE HOMOLOG* (*RBOHD;* At5g46910), *SENESCENCE-ASSOCIATED GENE113* (*SAG113*; At5g59220) and *ORESARA1* (*ORE1/NAC6,* At5g39610). Thesame *y*-axis scale was used for each gene across samples, with the scale maximum shown. Gene structures are depicted with the direction of transcription shown. (*E*) Number of differentially expressed genes (DEGs) (P_adj_<0.05) showing absolute differences independent of treatment responses, a comparison of Bay-0 and Lp2-6 read counts during the same treatment conditions. (*F*) Number of DEGs (P_adj_<0.05) showing accession × treatment interaction effects for each comparison.

**Fig. S3.**
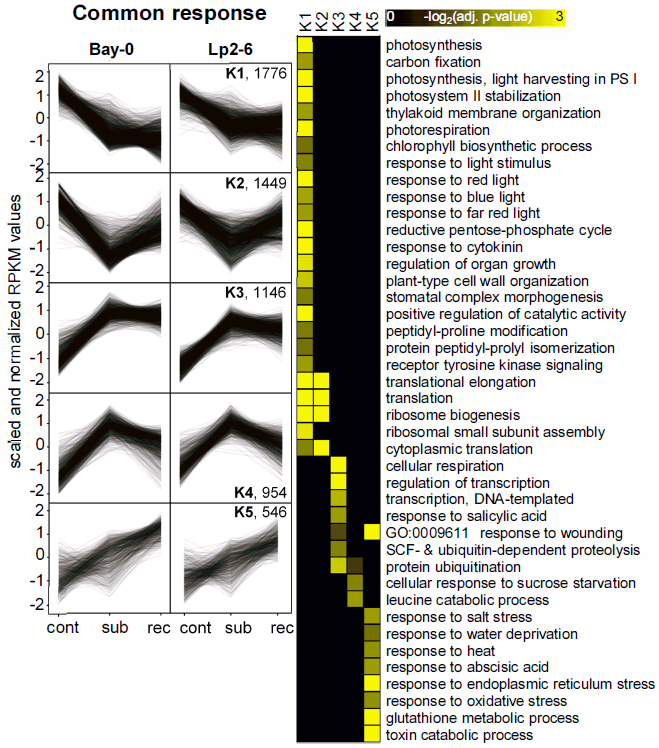
Common molecular processes in Bay-0 and Lp2-6 after submergence and 3 h of recovery. Fuzzy K-means plots visualize the regulation patterns of common response DEGs (P_adj_ <0.05) under control, submergence, and recovery. DEGs were individually plotted using RPKM values corrected for library size and library composition. GO analyses of identified clusters revealed associated biological 871 processes, where higher yellow color intensity indicates a stronger correlation between the genes 872 cluster and the GO term.

**Fig. S4.**
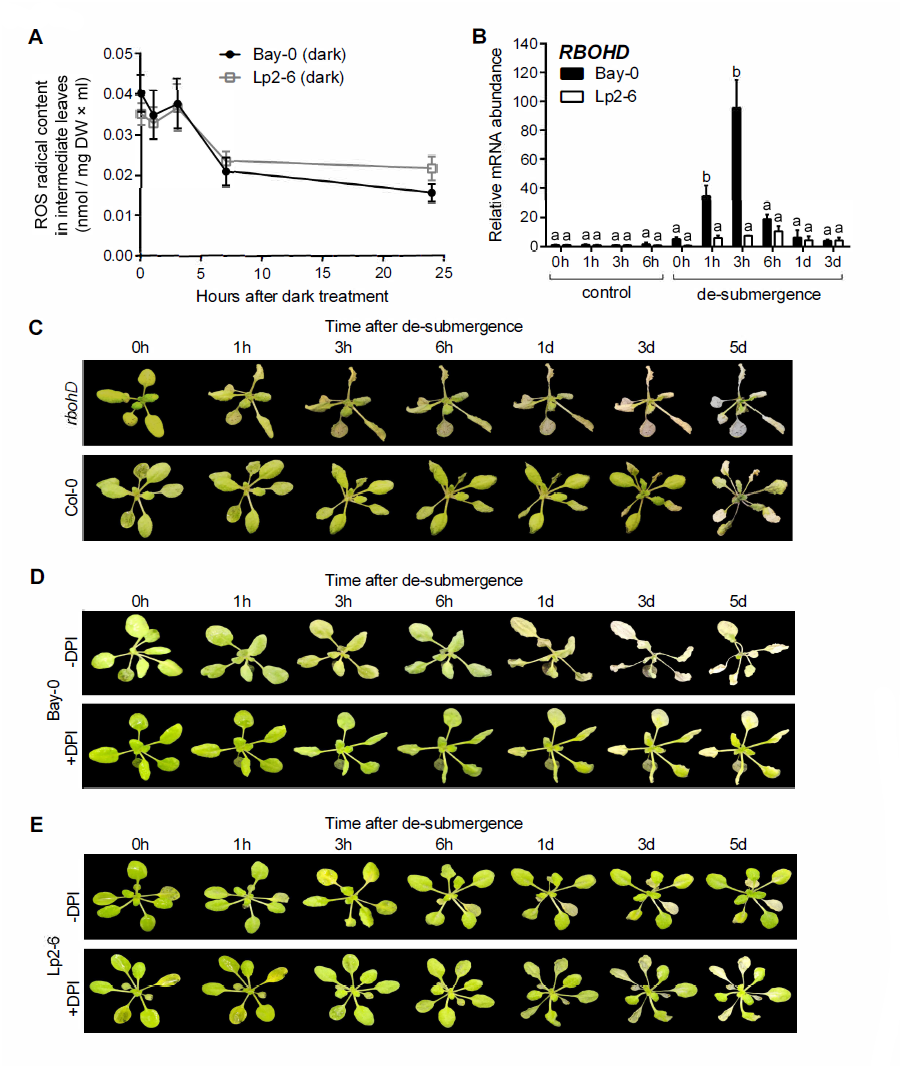
Controlled ROS production is required for recovery signaling. (*A*) Electron paramagnetic resonance (EPR) spectroscopy quantified ROS in Bay-0 and Lp2-6 intermediate leaves of control or recovering plants after 5 d of darkness (n=30). There was no significant difference (p<0.05) between the accessions at the specified time point. (*B*) Relative mRNA abundance of *RBOHD* measured by qRT-PCR in Bay-0 and Lp2-6 intermediate leaves following de-submergence after 5 d of submergence (n=3). Data represent mean ± SEM. Different letters represent significant difference (p<0.05, two-way ANOVA with Tukey’s multiple comparisons test). (*C*) Representative images of *rbohD* mutants and Col-0 wild-type plants recovering after 6 d of dark submergence. Representative images of recovering Bay-0 (*D*) and Lp2-6 (*E*) plants sprayed with 200 μM of the NADPH oxidase inhibitor DPI immediately upon de-submergence, following 5 d of submergence.

**Fig. S5.**
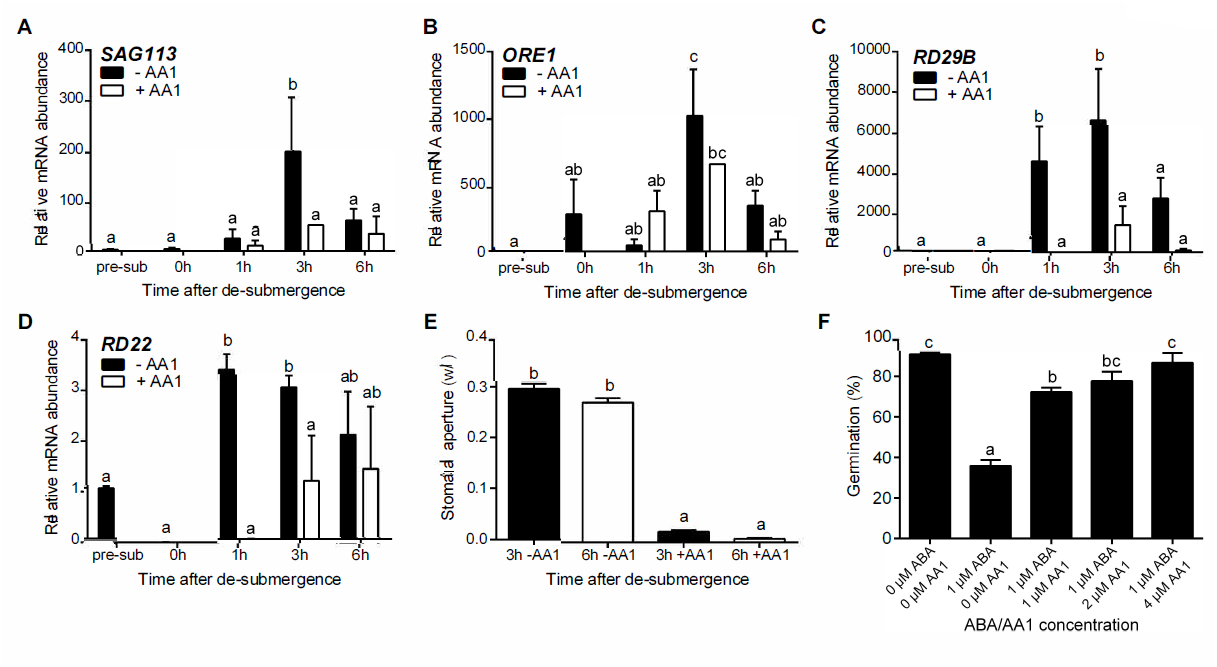
ABA regulation of *SAG113* and *ORE1*. Relative mRNA abundance of *SAG113* (*A*), *ORE1* (*B*), *RD29B* (*C*), and *RD22* (*D*) measured by qRT-PCR in intermediate leaves of Bay-0 before treatment (pre-sub), after 5 d of submergence (0 h) and subsequent recovery and treated with or without AA1 (n=3-4 biological replicates). (*E*) Stomatal aperture (based on width/length ratio) for Bay-0 intermediate leaves with or without 100 μM AA1 application upon de-submergence (n=300). (*F*) Seed germination rates of Col-0 on 1/2 MS medium with varying ABA and AA1 concentrations (n=5). Data represent mean ± SEM. Different letters represent significant difference (p<0.05, one-or two-way ANOVA with Tukey’s multiple comparisons test).

## Supplemental Movie

Time-lapse of a representative Bay-0 and Lp2-6 rosette recovering in normal growth conditions after 5 d of dark submergence.

## Supplemental Dataset

Log_2_FC, Benjamini-Hochberg-corrected P-values, and fuzzy K-means cluster number for all genes in the Ribo-seq dataset. Data is organized by 3 comparisons: submergence (plants submerged for 5 d in the dark compared to control plants), recovery (plants recovered for 3 h after de-submergence following 5 d of submergence compared to plants immediately de-submerged after 5 d of submergence), and combined response (plants recovered for 3 h compared to control plants).

**Table. S1.**
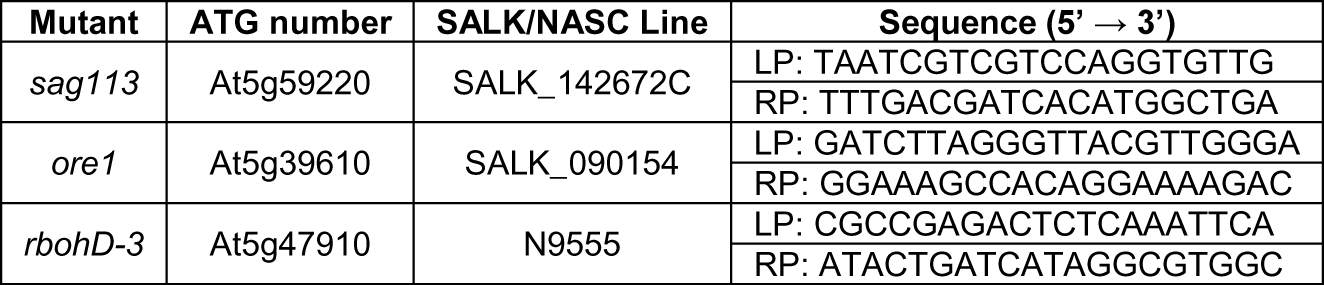
Primer sequences (5’ → 3’) for genotyping, indicated by the left primer (LP) and right primer (RP) of the insertion.

**Table. S2.**
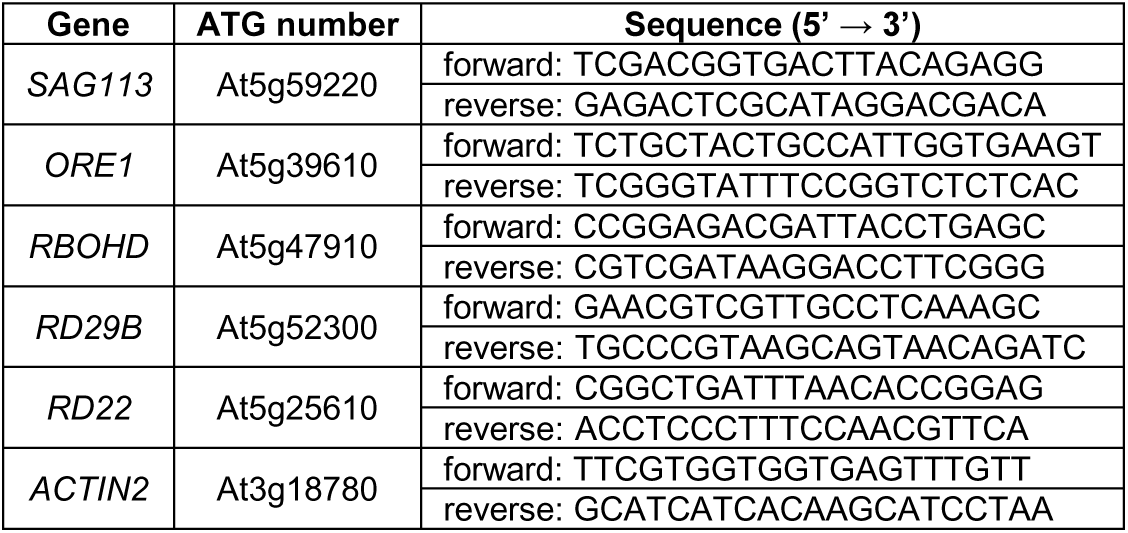
Primer sequences (5’ → 3’) for qRT-PCR.

## References

1. Bailey-Serres J, Lee SC, Brinton E (2012) Waterproofing crops: effective flooding survival strategies. Plant Physiol 160(4):1698–1709.

2. Voesenek LACJ, Bailey-Serres J (2015) Flood adaptive traits and processes: an overview. New Phytol 206(1):57–73.

3. Hirabayashi Y, et al. (2013) Global flood risk under climate change. Nat Clim Chang 3(9):816–821.

4. Jackson MB (1985) Ethylene and responses of plants to soil waterlogging and submergence. Annu Rev Plant Physiol 36(1):145–174.

5. Armstrong W (1980) Aeration in higher plants. Adv Bot Res 7:225–332.

6. Maurel C, Simonneau T, Sutka M (2010) The significance of roots as hydraulic rheostats. J Exp Bot 61(12):3191–3198.

7. Tamang BG, Magliozzi JO, Maroof MAS, Fukao T (2014) Physiological and transcriptomic characterization of submergence and reoxygenation responses in soybean seedlings. Plant Cell Environ 37(10):2350–2365.

8. Tsai K, Chou S, Shih M (2014) Ethylene plays an essential role in the recovery of Arabidopsis during post-anaerobiosis reoxygenation. Plant Cell Environ 37(10):2391–2405.

9. Fukao T, Yeung E, Bailey-Serres J (2011) The submergence tolerance regulator SUB1A mediates crosstalk between submergence and drought tolerance in rice. Plant Cell 23:412–427.

10. Ella ES, Kawano N, Ito O (2003) Importance of active oxygen-scavenging system in the recovery of rice seedlings after submergence. Plant Sci 165(1):85–93.

11. Biemelt S, Keetman U, Albrecht G (1998) Re-aeration following hypoxia or anoxia leads to activation of the antioxidative defense system in roots of wheat seedlings. Plant Physiol 116(2):651–658.

12. Monk LS, Fagerstedt K V, Crawford RM (1987) Superoxide dismutase as an anaerobic polypeptide: a key factor in recovery from oxygen deprivation in Iris pseudacorus? Plant Physiol 85(4):1016–1020.

13. Alpuerto JB, Hussain RMF, Fukao T (2016) The key regulator of submergence tolerance, *SUB1A*, promotes photosynthetic and metabolic recovery from submergence damage in rice leaves. Plant Cell Environ 39(3):672–684.

14. Khan MN, Komatsu S (2016) Characterization of post-flooding recovery-responsive enzymes in soybean root and hypocotyl. J Plant Biol 59(5):478–487.

15. Ye X, et al. (2016) Submergence causes similar carbohydrate starvation but faster post-stress recovery than darkness in *Alternanthera philoxeroides* plants. PLoS One 11(10):e0165193.

16. Yuan L, et al. (2017) Jasmonate regulates plant responses to reoxygenation through activation of antioxidant synthesis. Plant Physiol 173(3):1864–1880.

17. Tsai K, Lin C, Ting C, Shih M (2016) Ethylene-regulated glutamate dehydrogenase fine-tunes metabolism during anoxia-reoxygenation. Plant Physiol 172(3):1548–1562.

18. Ingolia NT, Ghaemmaghami S, Newman JRS, Weissman JS (2009) Genome-wide analysis in vivo of translation with nucleotide resolution using ribosome profiling. Science 324(5924):218–223.

19. Ingolia NT (2014) Ribosome profiling: new views of translation, from single codons to genome scale. Nat Rev Genet 15(3):205–213.

20. Juntawong P, Bailey-Serres J (2012) Dynamic light regulation of translation status in *Arabidopsis thaliana*. Front Plant Sci 3:66.

21. Branco-Price C, Kaiser KA, Jang CJH, Larive CK, Bailey-Serres J (2008) Selective mRNA translation coordinates energetic and metabolic adjustments to cellular oxygen deprivation and reoxygenation in *Arabidopsis thaliana*. Plant J 56(5):743–755.

22. Benina M, Ribeiro DM, Gechev TS, Mueller-Roeber B, Schippers JHM (2015) A cell type-specific view on the translation of mRNAs from ROS-responsive genes upon paraquat treatment of *Arabidopsis thaliana* leaves. Plant Cell Environ 38(2):349–363.

23. Cheng MC, et al. (2015) Increased glutathione contributes to stress tolerance and global translational changes in Arabidopsis. Plant J 83(5):926–939.

24. Sorenson R, Bailey-Serres J (2014) Selective mRNA translation tailors low oxygen energetics. Low-Oxygen Stress in Plants, eds van Dongen JT, Licausi F (Springer, Vienna), pp 95–115.

25. Dodge AD, Harris N (1970) The mode of action of paraquat and diquat. Biochem J 118(3):43P– 44P.

26. Bus JS, Aust SD, Gibson JE (1974) Superoxide- and singlet oxygen-catalyzed lipid peroxidation as a possible mechanism for paraquat (methyl viologen) toxicity. Biochem Biophys Res Commun 58(3):749–755.

27. Torres MA, Dangl JL, Jones JDG (2002) Arabidopsis gp91phox homologues AtrbohD and AtrbohF are required for accumulation of reactive oxygen intermediates in the plant defense response. Proc Natl Acad Sci USA 99(1):517–522.

28. Tissier AF, et al. (1999) Multiple independent defective suppressor-mutator transposon insertions in Arabidopsis: a tool for functional genomics. Plant Cell 11(10):1841–1852.

29. Vashisht D, et al. (2011) Natural variation of submergence tolerance among *Arabidopsis thaliana* accessions. New Phytol 190(2):299–310.

30. Lee SC, et al. (2011) Molecular characterization of the submergence response of the *Arabidopsis thaliana* ecotype Columbia. New Phytol 190(2):457–471.

31. Chang KN, et al. (2013) Temporal transcriptional response to ethylene gas drives growth hormone cross-regulation in Arabidopsis. Elife 2013(2):1–20.

32. Kim HJ, et al. (2014) Gene regulatory cascade of senescence-associated NAC transcription factors activated by ETHYLENE-INSENSITIVE2-mediated leaf senescence signalling in *Arabidopsis*. J Exp Bot 65(14):4023–4036.

33. Zhang K, Gan S (2012) An abscisic acid-AtNAP transcription factor-SAG113 protein phosphatase 2C regulatory chain for controlling dehydration in senescing Arabidopsis leaves. Plant Physiol 158(2):961–969.

34. Zhang K, Xia X, Zhang Y, Gan S (2012) An ABA-regulated and Golgi-localized protein phosphatase controls water loss during leaf senescence in Arabidopsis. Plant J 69(4):667–678.

35. He X, et al. (2005) AtNAC2, a transcription factor downstream of ethylene and auxin signaling pathways, is involved in salt stress response and lateral root development. Plant J 44(6):903–916.

36. Balazadeh S, et al. (2010) A gene regulatory network controlled by the NAC transcription factor ANAC092/AtNAC2/ORE1 during salt-promoted senescence. Plant J 62(2):250–264.

37. Qiu K, et al. (2015) EIN3 and ORE1 accelerate degreening during ethylene-mediated leaf senescence by directly activating chlorophyll catabolic genes in *Arabidopsis*. PLOS Genet 11(7):e1005399.

38. Buchanan-Wollaston V, et al. (2005) Comparative transcriptome analysis reveals significant differences in gene expression and signalling pathways between developmental and dark/starvation-induced senescence in Arabidopsis. Plant J 42(4):567–585.

39. Ye Y, et al. (2017) A novel chemical inhibitor of ABA signaling targets all ABA receptors. Plant Physiol 173(4):2356–2369.

40. Elstner EF, Osswald W (1994) Mechanisms of oxygen activation during plant stress. Proc R Soc Edinburgh 102:131–154.

41. Smirnoff N (1995) Antioxidant systems and plant response to the environment. Environment and Plant Metabolism: Flexibility and Acclimation (BIOS Scientific Publishers, UK).

42. Huang S, Van Aken O, Schwarzländer M, Belt K, Millar AH (2016) The roles of mitochondrial reactive oxygen species in cellular signaling and stress response in plants. Plant Physiol 171(3):1551–1559.

43. Baxter-Burrell A, Yang Z, Springer PS, Bailey-Serres J (2002) RopGAP4-dependent Rop GTPase rheostat control of *Arabidopsis* oxygen deprivation tolerance. Science 296(5575):2026–2028.

44. Pucciariello C, Parlanti S, Banti V, Novi G, Perata P (2012) Reactive oxygen species-driven transcription in Arabidopsis under oxygen deprivation. Plant Physiol 159(1):184–196.

45. Mustroph A, et al. (2009) Profiling translatomes of discrete cell populations resolves altered cellular priorities during hypoxia in *Arabidopsis*. Proc Natl Acad Sci USA 106(44):18843–18848.

46. Yao Y, et al. (2016) *ETHYLENE RESPONSE FACTOR 74 (ERF74*) plays an essential role in controlling a respiratory burst oxidase homolog D (RbohD)-dependent mechanism in response to different stresses in Arabidopsis. New Phytol 213(4):1667–1681.

47. Akman M, Kleine R, van Tienderen PH, Schranz EM (2017) Identification of the submergence tolerance QTL Come Quick Drowning1 (CQD1) in *Arabidopsis thaliana*. J Hered 108(3):308–317.

48. Setter TLT, et al. (2010) Desiccation of leaves after de-submergence is one cause for intolerance to complete submergence of the rice cultivar IR 42. Funct Plant Biol 37(11):1096–1104.

49. Shahzad Z, et al. (2016) A potassium-dependent oxygen sensing pathway regulates plant root hydraulics. Cell 167(1):87–98.e14.

50. Tanaka Y, et al. (2005) Ethylene inhibits abscisic acid-induced stomatal closure in Arabidopsis. Plant Physiol 138(4):2337–2343.

51. Voesenek LACJ, et al. (2003) De-submergence-induced ethylene production in *Rumex palustris*: regulation and ecophysiological significance. Plant J 33(2):341–352.

52. Porra RJ, Thompson WA, Kriedemann PE (1989) Determination of accurate extinction coefficients and simultaneous equations for assaying chlorophylls a and b extracted with four different solvents: verification of the concentration of chlorophyll standards by atomic absorption spectroscopy. Biochim Biophys Acta - Bioenerg 975(3):384–394.

53. Marsch-Martínez N, et al. (2013) An efficient flat-surface collar-free grafting method for *Arabidopsis thaliana* seedlings. Plant Methods 9(1):14.

54. Ingolia NT, Brar GA, Rouskin S, McGeachy AM, Weissman JS (2012) The ribosome profiling strategy for monitoring translation in vivo by deep sequencing of ribosome-protected mRNA fragments. Nat Protoc 7(8):1534–1550.

55. Juntawong P, Girke T, Bazin J, Bailey-Serres J (2014) Translational dynamics revealed by genome-wide profiling of ribosome footprints in *Arabidopsis*. Proc Natl Acad Sci USA 111(1):E203–E212.

56. Juntawong P, Hummel M, Bazin J, Bailey-Serres J (2015) Ribosome profiling: a tool for quantitative evaluation of dynamics in mRNA translation. Plant Functional Genomics (Humana Press, New York), pp 139–173.

57. Stewart RR, Bewley JD (1980) Lipid peroxidation associated with accelerated aging of soybean axes. Plant Physiol 65(2):245–8.

58. Zhang NX, et al. (2018) Phytophagy of omnivorous predator *Macrolophus pygmaeus* affects performance of herbivores through induced plant defences. Oecologia 186(1):101–113.

59. Livak KJ, Schmittgen TD (2001) Analysis of relative gene expression data using real-time quantitative PCR and the 2(-Delta Delta C(T)) method. Methods 25(4):402–408.

60. Backman TWH, Girke T (2016) systemPipeR: NGS workflow and report generation environment. BMC Bioinformatics 17:388.

61. Juntawong P, Bazin J, Hummel M, Bailey-Serres J, Girke T (2015) systemPipeR workflow for Ribo-seq and polyRibo-seq experiments. Available at www.bioconductor.org/. Accessed January 1, 2018.

62. Steffens B, Steffen-Heins A, Sauter M (2013) Reactive oxygen species mediate growth and death in submerged plants. Front Plant Sci 4:179.

